# Intervene: a tool for intersection and visualization of multiple gene or genomic region sets

**DOI:** 10.1101/109728

**Authors:** Aziz Khan, Anthony Mathelier

**Affiliations:** Centre for Molecular Medicine Norway (NCMM), Nordic EMBL Partnership, University of Oslo, 0349 Oslo, Norway; Department of Cancer Genetics, Institute for Cancer Research, Oslo University Hospital Radiumhospitalet, 0372 Oslo, Norway

**Keywords:** Visualization, Venn diagrams, UpSet plots, Heat maps, Genome analysis

## Abstract

**Background:** A common task for scientists relies on comparing lists of genes or genomic regions derived from high-throughput sequencing experiments. While several tools exist to intersect and visualize sets of genes, similar tools dedicated to the visualization of genomic region sets are currently limited.

**Results:** To address this gap, we have developed the Intervene tool, which provides an easy and automated interface for the effective intersection and visualization of genomic region or list sets, thus facilitating their analysis and interpretation. Intervene contains three modules: *venn* to generate Venn diagrams of up to six sets, *upset* to generate UpSet plots of multiple sets, and *pairwise* to compute and visualize intersections of multiple sets as clustered heat maps. Intervene, and its interactive web ShinyApp companion, generate publication-quality figures for the interpretation of genomic region and list sets.

**Conclusions:** Intervene and its web application companion provide an easy command line, and an interactive web interface to compute intersections of multiple genomic and list sets. They also have the capacity to plot intersections using easy-to-interpret visual approaches. Intervene is developed and designed to meet the needs of both computer scientists and biologists. The source code is freely available at https://bitbucket.org/CBGR/intervene, with the web application available at https://asntech.shinyapps.io/intervene.

## Background

Effective visualization of transcriptomic, genomic, and epigenomic data generated by next-generation sequencing-based high-throughput assays have become an area of great interest. The majority of the data sets generated by such assays are lists of genes or variants, and genomic region sets. The genomic region sets represent genomic locations for specific features, such as transcription factor – DNA interactions, transcription start sites, histone modifications, and DNase hypersensitivity sites. A common task in the interpretation of these features is to find similarities, differences, and enrichments between such sets, which come from different samples, experimental conditions, or cell and tissue types.

Classically, the intersection or overlap between different sets, such as gene lists, is represented by Venn diagrams [1] or Edwards-Venn [2]. If the number of sets exceeds four, such diagrams become complex and difficult to interpret. The key challenge is that there are *2^n^* combinations to visually represent when considering *n* sets. An alternative approach, the UpSet plots, was introduced to depict the intersection of more than three sets [3]. The advantage of UpSet plots is their capacity to rank the intersections and alternatively hide combinations without intersection, which is not possible using a Venn diagram. However, with a large number of sets, UpSet plots become an ineffective way of illustrating set intersections. To visualize a large number of sets, one can represent pairwise intersections using a clustered heat map as suggested in [4].

There are several web applications and R packages available to compute intersection and visualization of up-to six list sets by using Venn diagrams. Although tools exist to perform genomic region set intersections [5-7], there is a limited number of tools available to visualize them [5, 6]. To our knowledge no tool exists to generate UpSet plots for genomic region sets. As a consequence, there is a great need for integrative tools to compute and visualize intersection of multiple sets of both genomic regions and gene/list sets.

To address this need, we developed Intervene, an easy-to-use command line tool to compute and visualize intersections of genomic regions with Venn diagrams, UpSet plots, or clustered heat maps. Moreover, we provide an interactive web application companion to upload list sets or the output of Intervene to further customize plots.

## Implementation

Intervene comes as a command line tool, along with an interactive Shiny web application to customize the visual representation of intersections. The command line tool is implemented in Python (version 2.7) and R programming language (version 3.3.2). The build also works with Python versions 3.4, 3.5, and 3.6. The accompanying web interface is developed using Shiny (version 1.0.0), a web application framework for R. Intervene uses pybedtools [6] to perform genomic region set intersections and Seaborn (https://seaborn.pydata.org/), Matplotlib [7], UpSetR [3], and Corrplot [8] to generate figures. The web application uses the R package Venerable [9] for different types of Venn diagrams, UpSetR for UpSet plots, and heatmap.2 and Corrplot for pairwise intersection clustered heat maps.

Intervene can be installed by using *pip install intervene* or using the source code available on bitbucket https://bitbucket.org/CBGR/intervene. The tool has been tested on Linux and MAC systems. The Shiny web application is hosted with shinyapps.io by RStudio, and is compatible with all modern web browsers. A detailed documentation including installation instructions and how to use the tool is provided in **Additional file 1** and is available at http://intervene.readthedocs.io.

## Results

### An integrated tool for effective visualization of multiple set intersections

As visualization of sets and their intersections is becoming more and more challenging due to the increasing number of generated data sets, there is a strong need to have an integrated tool to compute and visualize intersections effectively. To address this challenge, we have developed Intervene, which is composed of three different modules, accessible through the subcommands *venn, upset*, and *pairwise*. A detailed sketch of input types, subcommands, and the web application utility is provided in **Figure 1.**

**Figure 1:**
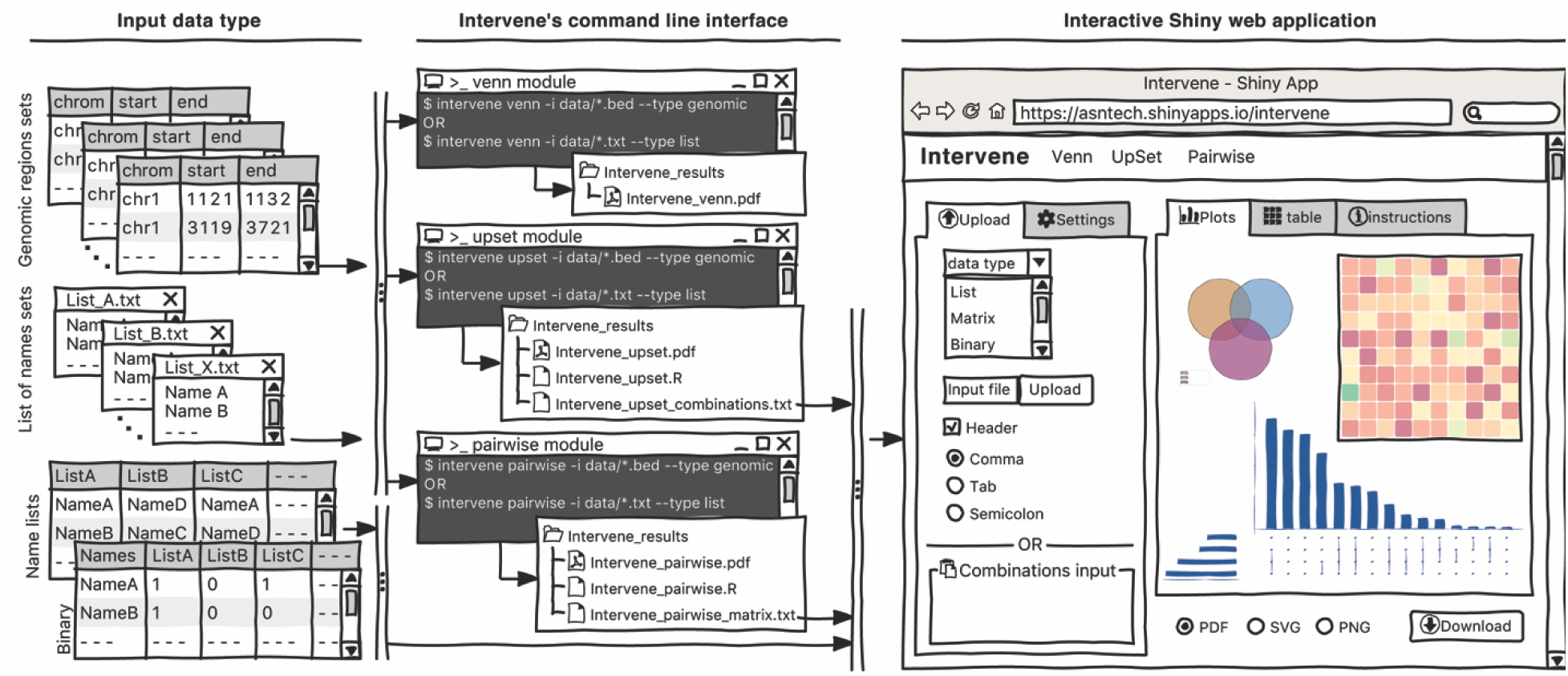
A sketch of Intervene’s command line interface and Shiny web application with input data type.

Intervene provides flexibility to the user to choose figure colors, label text, size, resolution, and type to make them publication-standard quality. To access the help function within any module, the user can type *intervene <subcommand> ‐‐help* on the command line. Furthermore, Intervene produces results as text files, which can be easily imported to the web application for interactive visualization and customization of plots (see section “An interactive web application”).

### Venn diagrams module

Venn diagrams are the classical approach to show intersections between sets. There are several web-based applications and R packages available to visualize intersections of up-to six list sets in classical Venn, Euler, or Edward’s diagrams [10–15]. However, a very limited number of tools are available to visualize genomic region intersections using classical Venn diagrams [5, 6].

Intervene provides up-to six-way classical Venn diagrams for gene lists or genomic region sets. The associated web interface can also be used to compute the intersection of multiple gene sets, and visualize it using different flavors of weighted and unweighted Venn and Euler diagrams. These different types include: classical Venn diagrams (up-to five sets), Chow-Ruskey (up-to five sets), Edwards’ diagrams (up-to five sets), and Battle (up-to nine sets).

As an example, one might be interested to calculate the number of overlapping ChIP-seq (chromatin immunoprecipitation followed by sequencing) peaks between different types of histone modification marks (H3K27ac, H3K4me3, and H3K27me3) in human embryonic stem cells (hESC) [16] (**Figure 2a**, can be generated with the command *intervene venn ‐‐test*).

**Figure 2:**
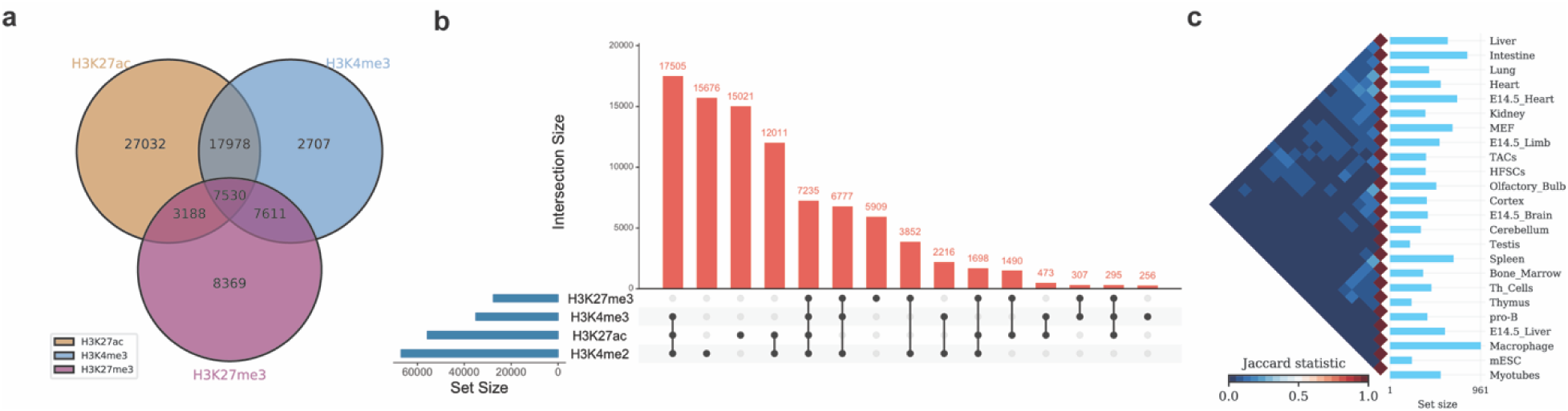
Example of Intervene’s command line interface outputs. (a)A three-way Venn diagram of ChIP-seq peaks of histone modifications (H3K27ac, H3Kme3 and H3K27me3) in hESC obtained from ENCODE (b) UpSet plot of the intersection of four histone modification peaks in hESC (c) A heatmap of pairwise intersections terms of Jaccard statistics of super-enhancers in 24 mouse cell and tissue types downloaded from dbSUPER.

### UpSet plots module

When the number of sets exceeds four, Venn diagrams become difficult to read and interpret. An alternative and more effective approach is to use UpSet plots to visualize the intersections. An R package and an interactive web-based tool are available at http://vcg.github.io/upset to visualize multiple list sets. However, to our knowledge, there is no tool available to draw the UpSet plots for genomic region set intersections. Intervene’s *upset* subcommand can be used to visualize the intersection of multiple genomic region sets using UpSet plots.

As an example, we show the intersections of ChIP-seq peaks for histone modifications (H3K27ac, H3K4me3, H3K27me3 and H3K4me2) in hESC using an UpSet plot, where interactions were ranked by frequency (**Figure 2b,** can be generated with the command *intervene upset ‐‐test*). This plot is easier to understand than the four-way Venn diagram (**Additional file 1**).

### Pairwise intersection heat maps module

With an increasing number of data sets, visualizing all possible intersections becomes unfeasible by using Venn diagrams or UpSet plots. One possibility is to compute pairwise intersections and plot-associated metrics as a clustered heat map. Intervene’s *pairwise* module provides several metrics to assess intersections, including number of overlaps, fraction of overlap, Jaccard statistics, Fisher’s exact test, and distribution of relative distances. Moreover, the user can choose from different styles of heat maps and clustering approaches.

As an example, we obtained the genomic regions of super enhancers in 24 mouse cell type and tissues from dbSUPER [17] and computed the pairwise intersections in terms of Jaccard statistics (**Figure 2c**). The triangular heat map shows the pairwise Jaccard index, which is between 0 and 1, where 0 means no overlap and 1 means full overlap. The bar plot shows the number of regions in each cell-type or tissue. This plot can be generated using the command *intervene pairwise ‐‐test*).

### An interactive web application

Intervene comes with a web application companion to further explore and filter the results in an interactive way. Indeed, intersections between large data sets can be computed locally using Intervene’s command line interface, then the output files can be uploaded to the ShinyApp for further exploration and customization of the figures (**Figure 1**). The ShinyApp web interface takes four types of inputs: (i) a text/csv file where each column represents a set, (ii) a binary representation of intersections, (iii) a pairwise matrix of intersections, and (iv) a matrix of overlap counts. The web application provides several easy and intuitive customization options for responsive adjustments of the figures (**Figures 1** and **3**). Users can change colors, fonts and plot sizes, change labels, and select and deselect specific sets. These customized and publication-ready figures can be downloaded in PDF, SVG, TIFF, and PNG formats. The pairwise modules also provides three types of correlation coefficients and hierarchical clustering with eight clustering methods and four distance measurement methods. It further provides interactive features to explore data values; this is done by hovering the mouse cursor over each heat map cell, or by using a searchable and sortable data table. The data table can be downloaded as a CSV file and interactive heat maps can be downloaded as HTML. The Shiny-based web application is freely available at https://asntech.shinyapps.io/intervene.

**Figure 3:**
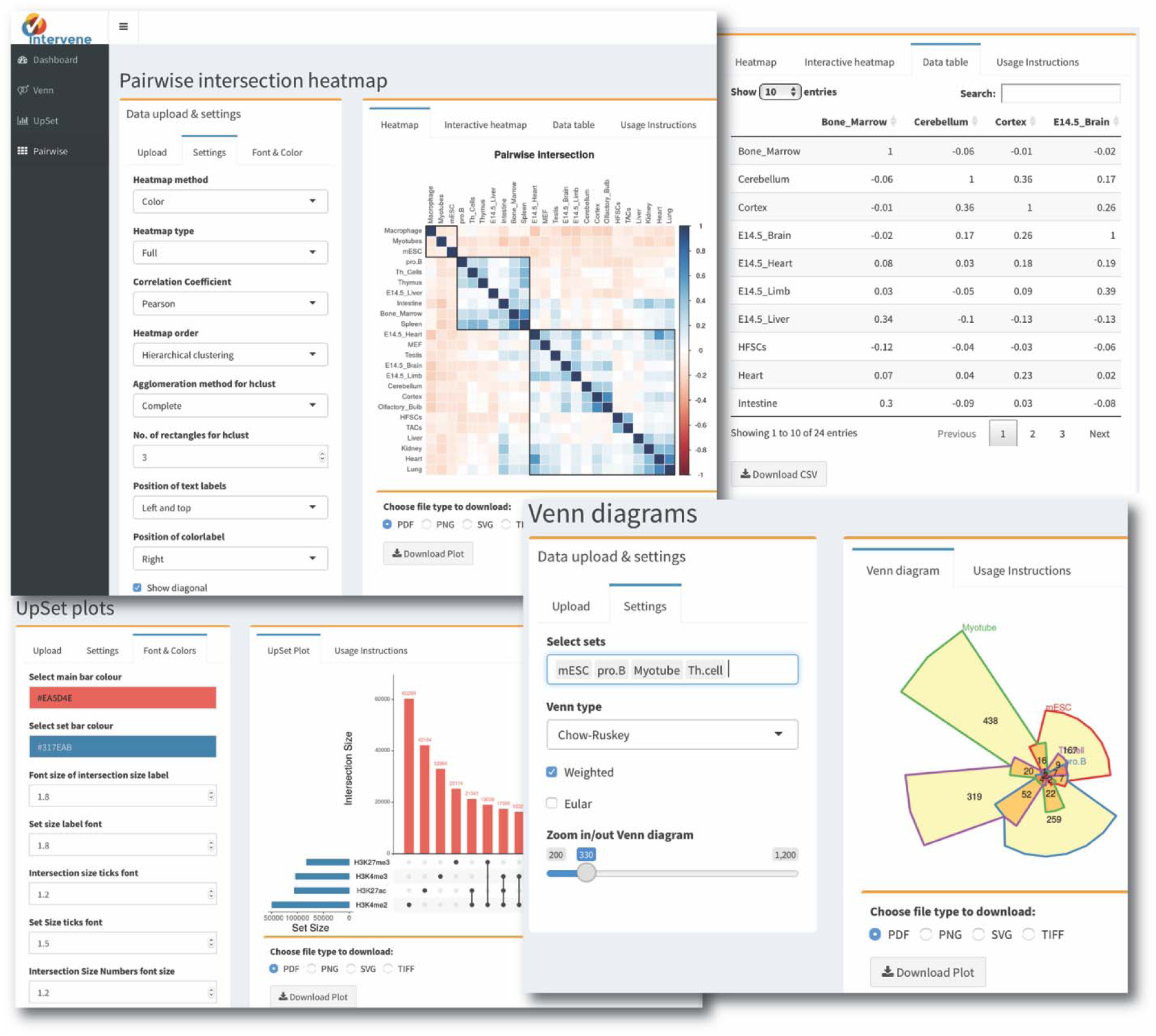
Screenshot of web application user interface.

### Case study: highlighting co-binding factors in the MCF-7 cell line

Transcription factors (TFs) are key proteins regulating transcription through their cooperative binding to the DNA [18, 19]. To highlight Intervene’s capabilities, we used the command-line tool and its ShinyApp companion to predict and visualize cooperative interactions between TFs at cis-regulatory regions in the MCF-7 breast cancer cell line. Specifically, we considered (i) TF binding regions derived from uniformly processed TF ChIP-seq experiments compiled in the ReMap database [20] and (ii) promoter and enhancer regions predicted by chromHMM [21] from histone modifications and regulatory factors ChIP-seq [22]. The pairwise module of Intervene was used to compute the fraction of overlap between all pairs of ChIP-seq data sets and regulatory regions. The output matrix was provided to the ShinyApp to compute Spearman correlations of the computed values and to generate the corresponding clustering heat map (default parameters; Figure 4). The largest cluster (green cluster) was composed of the three key cooperative TFs involved in oestrogen-positive breast cancers: ESR1, FOXA1, and GATA3. They were clustered with enhancer regions where they have been shown to interact [23]. The cluster highlights potential TF cooperators: ARNT, AHR, GREB1, and TLE3. Promoter regions were found in the second largest cluster (red cluster), along with CTCF, STAG1, and RAD21, which are known to orchestrate chromatin architecture in human cells [24]. The last cluster was principally composed by TFAP2C data sets. Taken together, Intervene visually highlighted the cooperation of different sets TFs at MCF-7 promoters and enhancers, in agreement with the literature.

**Figure 4:**
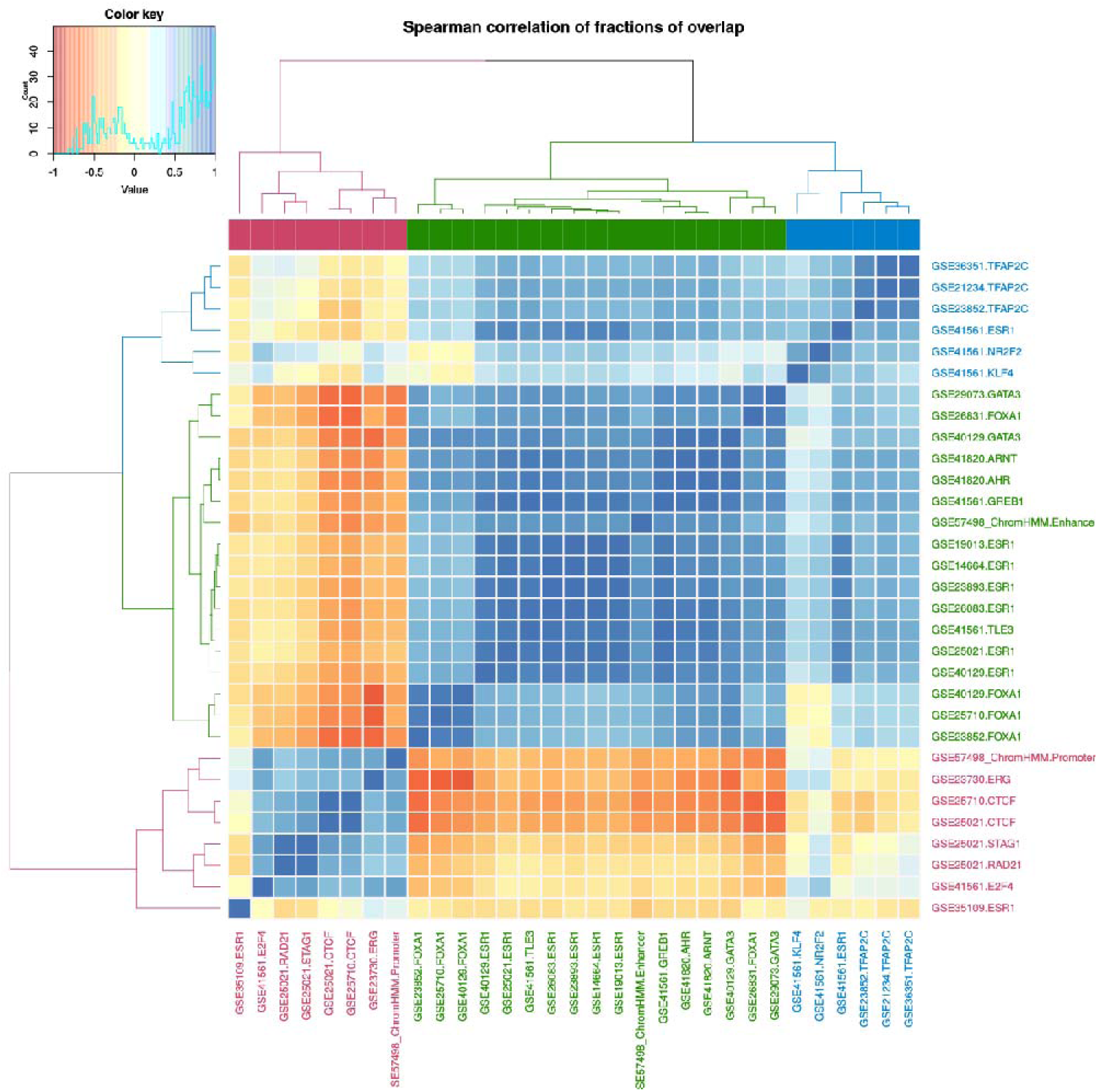
MCF-7 cluster heat map. Cluster heat map of the Spearman correlations of fractions of overlap between TF ChIP-seq data sets and regulatory regions in MCF-7. Three clusters (red, green, and blue) are highlighted.

## Discussion

A comparative analysis of different tools to compute and visualize intersections as Venn diagrams, UpSet plots, and pairwise heat maps is provided in Table 1. Most of the tools available currently can only draw Venn diagrams for up-to six list sets. Intervene provides Venn diagrams, UpSet plots, and pairwise heat maps for both list sets and genomic region sets. To the best of our knowledge, it is the only tool available to draw UpSet plots for the intersections of genomic region sets. Intervene is the first of its kind to allow for the computation and visualization of intersections between multiple genomic region and list sets with three different approaches.

**Table 1:** Comparison of Intervene with currently available tools to draw Venn diagrams, UpSet plots and pairwise heatmaps.

| Application | Venn plot types | Upset plot | Pairwise heat map | Weighted venn | Application type | Input type | No. Of inputs | Output type |
| --- | --- | --- | --- | --- | --- | --- | --- | --- |
| VennDiagramWeb [11] | Classical Venn, Euler | ✗ | ✗ | ✗ | web app | Lists | 5 | TIFF, SVG, PNG, R objects |
| VennPainter [13] | Classical Venn, Edwards, Nested Venn | ✗ | ✗ | ✗ | Stand-alone | Lists | 8 | SVG, text |
| Vennture [14] | Edwards | ✗ | ✗ | ✗ | Stand-alone | Lists | 6 | PowerPoint, Excel |
| BioVenn [10] | Classical Venn | ✗ | ✗ | ✓ | web app | Lists | 3 | SVG, PNG |
| jVenn [12] | Classical Venn, Edwards | ✗ | ✗ | ✗ | web app | Lists | 6 | PNG and SVG |
| InteractiVenn [15] | Edwards | ✗ | ✗ | ✗ | web app | Lists | 6 | SVG, PNG, text |
| UpSet [3] | ✗ | ✓ | ✗ | ✗ | web app, R package | Lists, binary, counts | Multiple | PDF, PNG |
| ChippeakAnno [5] | Classical Venn | ✗ | ✗ | ✗ | R package | Genomic regions | 5 | PDF, SVG, PNG |
| Pybedtools [6] | Classical Venn | ✗ | Matrix only | ✓ | command line | Genomic regions | 3 for Venn, multiple for pairwise | PDF, SVG, PNG |
| Intervene | Classical Venn, Euler, Edwards, Chow-Ruskey, Square, Battle | ✓ | ✓ | ✓ | command line, web app | Genomic regions, lists, binary, counts | 6 for Venn, multiple for upset and pairwise | PDF, SVG, PNG, TIFF, R objects, text |

In the near future, Intervene will be integrated to the Galaxy Tool Shed to be easily installed to any Galaxy instance with one click. We plan to develop a dedicated web application allowing users to upload genomic region sets for intersections and visualization.

## Conclusion

We described Intervene as an integrated tool that provides an easy and automated interface for intersection, and effective visualization of genomic region and list sets. To our knowledge, Intervene is the first tool to provide three types of visualization approaches for multiple sets of gene or genomic intervals. The three modules are developed to overcome the situations where the number of sets is large. Intervene and its web application companion are developed and designed to fit the needs of a wide range of scientists.

## Abbreviations

hESCs: Human embryonic stem cells
SEs: Super-enhancers
ENCODE: The Encyclopedia of DNA Elements
ChIP-seq: Chromatin immunoprecipitation followed by sequencing
TFs: Transcription factors

## Declarations

### Ethics approval and consent to participate

Not applicable.

### Consent for publication

Not applicable.

### Availability of data and materials

The source code of Intervene and test data are freely available at https://bitbucket.org/CBGR/intervene and a detailed documentation can be found at http://intervene.readthedocs.io. An interactive Shiny App is available at https://asntech.shinyapps.io/intervene

### Author’s contributions

AK conceived the project. AK and AM designed the tool. AM supervised the project. AK implemented both Intervene and the Shiny web application. AK wrote the manuscript draft and AM revised it. All authors read and approved the manuscript.

## Acknowledgements

We thank Marius Gheorghe and Dimitris Polychronopoulos for their useful suggestions and testing the tool, and Annabel Darby for providing suggestions on the manuscript text.

## References

1. Venn J: On the diagrammatic and mechanical representation of propositions and reasonings. Philos Mag J Sci 1880, 10:1–18.

2. Edwards AWF: Cogwheels of the Mind: The Story of Venn Diagrams. JHU Press; 2004.

3. Lex A, Gehlenborg N, Strobelt H, Vuillemot R, Pfister H: UpSet: Visualization of intersecting sets. IEEE Trans Vis Comput Graph 2014, 20:1983–1992.

4. Lex A, Gehlenborg N: Points of view: Sets and intersections. Nat Meth 2014, 11:779.

5. Zhu LJ, Gazin C, Lawson ND, Pagès H, Lin SM, Lapointe DS, Green MR: ChIPpeakAnno: a Bioconductor package to annotate ChIP-seq and ChIP-chip data. BMC Bioinformatics 2010, 11:237.

6. Dale RK, Pedersen BS, Quinlan AR: Pybedtools: A flexible Python library for manipulating genomic datasets and annotations. Bioinformatics 2011, 27:3423–3424.

7. Hunter JD: Matplotlib: A 2D graphics environment. Comput Sci Eng 2007, 9:99–104.

8. Wei T, Simko V: corrplot: Visualization of a Correlation Matrix. R package version 0.77. https://CRAN.R-project.org/package=corrplot.

9. Swinton J: Vennerable: Venn and Euler area-proportional diagrams. R package version 3.1.0.9000. https://github.com/js229/Vennerable

10. Hulsen T, de Vlieg J, Alkema W: BioVenn – a web application for the comparison and visualization of biological lists using area-proportional Venn diagrams. BMC Genomics 2008, 9:488.

11. Lam F, Lalansingh CM, Babaran HE, Wang Z, Prokopec SD, Fox NS, Boutros PC: VennDiagramWeb: a web application for the generation of highly customizable Venn and Euler diagrams. BMC Bioinformatics 2016, 17:401.

12. Bardou P, Mariette J, Escudié F, Djemiel C, Klopp C: jvenn: an interactive Venn diagram viewer. BMC Bioinformatics 2014, 15:293.

13. Lin G, Chai J, Yuan S, Mai C, Cai L, Murphy RW, Zhou W, Luo J: VennPainter: A Tool for the Comparison and Identification of Candidate Genes Based on Venn Diagrams. PLoS One 2016, 11:e0154315.

14. Martin B, Chadwick W, Yi T, Park S-S, Lu D, Ni B, Gadkaree S, Farhang K, Becker KG, Maudsley S: VENNTURE–A Novel Venn Diagram Investigational Tool for Multiple Pharmacological Dataset Analysis. PLoS One 2012, 7:e36911.

15. Heberle H, Meirelles GV, da Silva FR, Telles GP, Minghim R: InteractiVenn: a web-based tool for the analysis of sets through Venn diagrams. BMC Bioinformatics 2015, 16:169.

16. Dunham I, Kundaje A, Aldred SF, Collins PJ, Davis C a., Doyle F, Epstein CB, Frietze S, Harrow J, Kaul R, Khatun J, Lajoie BR, Landt SG, Lee B-K, Pauli F, Rosenbloom KR, Sabo P, Safi A, Sanyal A, Shoresh N, Simon JM, Song L, Trinklein ND, Altshuler RC, Birney E, Brown JB, Cheng C, Djebali S, Dong X, Ernst J, et al.: An integrated encyclopedia of DNA elements in the human genome. Nature 2012, 489:57–74.

17. Khan A, Zhang X: dbSUPER: a database of super-enhancers in mouse and human genome. Nucleic Acids Res 2016, 44(Database issue):D164–D171.

18. Papp B, Sabri S, Ernst J, Plath K: Cooperative Binding of Transcription Factors Orchestrates Reprogramming. Cell 2017:1–18.

19. Spitz F, Furlong EEM: Transcription factors: from enhancer binding to developmental control. Nat Rev Genet 2012, 13:613–26.

20. Griffon A, Barbier Q, Dalino J, Van Helden J, Spicuglia S, Ballester B: Integrative analysis of public ChIP-seq experiments reveals a complex multi-cell regulatory landscape. Nucleic Acids Res 2015, 43:1–14.

21. Ernst J, Kellis M: ChromHMM: automating chromatin-state discovery and characterization. Nat Methods 2012, 9:215–216.

22. Taberlay PC, Statham AL, Kelly TK, Clark SJ, Jones PA: Reconfiguration of nucleosome-depleted regions at distal regulatory elements accompanies DNA methylation of enhancers and insulators in cancer. Genome Res 2014, 24:1421–1432.

23. Theodorou V, Stark R, Menon S, Carroll JS: GATA3 acts upstream of FOXA1 in mediating ESR1 binding by shaping enhancer accessibility. Genome Res 2013, 23:12–22.

24. Zuin J, Dixon JR, van der Reijden MIJA, Ye Z, Kolovos P, Brouwer RWW, van de Corput MPC, van de Werken HJG, Knoch TA, van IJcken WFJ, Grosveld FG, Ren B, Wendt KS: Cohesin and CTCF differentially affect chromatin architecture and gene expression in human cells. Proc Natl Acad Sci USA 2014, 111:996–1001.

